# Diclofenac and other Non-Steroidal Anti-Inflammatory Drugs (NSAIDs) are Competitive Antagonists of the human P2X3 Receptor

**DOI:** 10.1101/2022.12.19.520978

**Authors:** Laura Grohs, Linhan Cheng, Saskia Cönen, Bassam G. Haddad, Astrid Obrecht, Idil Toklucu, Lisa Ernst, Jannis Körner, Günther Schmalzing, Angelika Lampert, Jan-Philipp Machtens, Ralf Hausmann

## Abstract

The P2X3 receptor (P2X3R), an ATP-gated non-selective cation channel of the P2X receptor family, is expressed in sensory neurons and involved in nociception. P2X3R inhibition was shown to reduce chronic and neuropathic pain. In a previous screening of 2000 approved drugs, natural products and bioactive substances, various non-steroidal anti-inflammatory drugs (NSAIDs) were found to inhibit P2X3R-mediated currents. To investigate whether the inhibition of P2X receptors contributes to the analgesic effect of NSAIDs, we characterized the potency and selectivity of various NSAIDs at P2X3R and other P2XR subtypes using two-electrode voltage clamp electrophysiology. We identified diclofenac as a hP2X3R and hP2X2/3R antagonist with micromolar potency (with IC_50_ values of 138.2 µM and 76.7 µM, respectively). A weaker inhibition of hP2X1R, hP2X4R and hP2X7R by diclofenac was determined. Flufenamic acid (FFA) proved to inhibit hP2X3R, rP2X3R and hP2X7R (IC_50_ values of 221µM, 264.1µM and ∼ 900µM, respectively), questioning its widespread use as a nonselective ion channel blocker, when P2XR-mediated currents are under study.

Inhibition of the hP2X3R or hP2X2/3R by diclofenac could be overcome by prolonged ATP-application or increasing concentrations of the agonist α,β-meATP, respectively, indicating competition of diclofenac and the agonists. Molecular dynamics simulation showed that diclofenac largely overlaps with ATP bound to the open state of the hP2X3R. Our results strongly support a competitive antagonism through which diclofenac, by interacting with residues of the ATP-binding site, left flipper, and dorsal fin domains inhibits gating of P2X3R by conformational fixation of the left flipper and dorsal fin domains.

In summary, we demonstrate the inhibition of the human P2X3 receptor by various NSAIDs. Diclofenac proved to be the most effective antagonist with a strong inhibition of hP2X3R and hP2X2/3R and a weaker inhibition of hP2X1R, hP2X4R and hP2X7R. Considering their involvement in nociception, inhibition of hP2X3R and hP2X2/3R by micromolar concentrations of diclofenac may contribute to the analgesic effect as well as the side effect of taste disturbances of diclofenac and represent an additional mode of action besides the well-known high potency COX inhibition.

## 1 Introduction

P2X receptors (P2XR) constitute a family of non-selective cation channels gated by extracellular ATP (North and Barnard, 1997). Seven different subtypes (P2X1-7) can assemble to homo- or heterotrimers (Nicke et al., 1998, North, 2002).

For the P2X3R, which is expressed in sensory neurons (Chen et al., 1995), a crucial role in nociception has been demonstrated (Burnstock, 2016). P2X3-deficient mice exhibit an attenuated pain behavior after injection of ATP into the hind paw compared to wild-type mice (Cockayne et al., 2000), whereas the response to acute mechanical pain stimuli remains unchanged (Souslova et al., 2000). Accordingly, the pharmacological inhibition of P2X3R has been shown to effectively reduce chronic or neuropathic pain in rodents (Jarvis et al., 2002). Recently, the modulator TMEM163, a 289 amino acid transmembrane protein, was identified to be required for full function of the neuronal P2X3R- and P2X4R and pain-related ATP-evoked behavior in mice (Salm et al., 2020). In addition to P2X3R, an involvement in nociception could also be assigned to heterotrimeric P2X2/3R, P2X4R and P2X7R (Chessell et al., 2005, Cockayne et al., 2005, Tsuda et al., 2009). All of these seem to be more relevant for the development of neuropathic or inflammatory pain than for acute nociception (Chessell et al., 2005, Tsuda et al., 2009).

The important role of the P2X3R in nociception makes the P2X3R a promising target for the development of new analgesics (North and Jarvis, 2013). However, to this day none of the developed antagonists has been approved for clinical use as an analgesic, even if Gefapixant (formerly AF-219) is approved as an anti-cough agent in Japan (details are given below). One of the first potent and selective P2X3R (and P2X2/3R) antagonists was A-317491, which successfully reduced chronic pain in rodent models (Jarvis et al., 2002), but showed insufficient distribution into the central nervous system (Sharp et al., 2006). Several other P2X3R antagonists have been developed as clinical candidates, such as AF-219/gefapixant, BAY-1817080/eliapixant, BLU-5937, MK-3901 or S-600918/sivopixant (Niimi et al., 2022, Spinaci et al., 2021). The availability of the crystal structures of the human P2X3R (Mansoor et al., 2016), together with cryo-EM techniques is ideally suited to facilitate structure-based drug design for P2X3Rs by revealing and characterizing novel ligand-binding sites (Oken et al., 2022).

The most advanced is the development of gefapixant, a P2X3R and P2X2/3R antagonist, which effectively reduced chronic cough caused by hypersensitivity of the cough reflex in phase 2 and 3 trials (Abdulqawi et al., 2015, Marucci et al., 2019). However, a taste disturbance was described as a side effect by all patients (Abdulqawi et al., 2015). Gefapixant as a first-in-class, non-narcotic, selective P2X3 receptor antagonist was recently approved for marketing in Japan as treatment for refractory or unexplained chronic cough (Markham, 2022). Another promising substance, BLU-5937, was able to effectively reduce chronic cough in animal models without altering taste sensation, possibly due to its considerably higher selectivity for P2X3R versus P2X2/3R (Garceau and Chauret, 2019). BLU-5937 is now part of a phase 2 study for the treatment of chronic cough (Marucci et al., 2019). Also, sivopixant was shown to reduce objective cough frequency and improved health-related quality of life, with a low incidence of taste disturbance, among patients with refractory or unexplained chronic cough in a phase 2a trial (Niimi et al., 2022). Eliapixant showed in its first-in-human study a favorable tolerability with no taste-related adverse events, and in a phase 1/2a study eliapixant showed reduced cough frequency and severity and was well tolerated with acceptable rates of taste-related events (Klein et al., 2022, Morice et al., 2014).

In light of the promising role of P2X3R antagonists for the treatment of pain and refractory cough, as well as the high likelihood of taste disturbances caused by not fully selective P2X3R antagonists (against heteromeric P2X2/3R), it appears interesting to investigate whether already approved drugs do affect the P2X3R mediated responses. For this purpose, a screening of 2000 approved drugs, natural products and bioactive substances was performed in a previous study of our group. In this screening, aurintricarboxylic acid (ATA) was identified as a potent P2X1R and P2X3R antagonist (Obrecht et al., 2019). An inhibitory effect could also be demonstrated for other drugs. These included various non-steroidal anti-inflammatory drugs (NSAIDs) and diclofenac showed the highest inhibitory effect of the screened NSAIDs. The analgesic, antipyretic and anti-inflammatory effect of NSAIDs is generally described to result from the inhibition of prostaglandin synthesis by inhibiting the cyclooxygenases COX-1 and COX-2 (Vane, 1971). Most NSAIDs constitute reversible, competitive blockers of the enzyme Cyclooxygenase (COX), while acetylsalicylic acid (Aspirin^®^) can cause an irreversible inactivation of COX through acetylation of Serine 530 (DeWitt et al., 1990, Rome and Lands, 1975).

Considering the involvement of P2X3R in nociception, it is conceivable that inhibition of P2X3R by NSAIDs represents an additional mode of action besides COX inhibition. To investigate whether the inhibition of P2XR contributes to the analgesic effect of NSAIDs, we determined the potency and selectivity of various NSAIDs at P2X3R and other P2XR subtypes using two-electrode voltage clamp (TEVC) electrophysiology. The investigated NSAIDs included diclofenac, ibuprofen, flunixin, meclofenamic acid, naproxen and flufenamic acid (FFA). The latter additionally plays an important role in research as a nonselective ion channel blocker (Guinamard et al., 2013).

In the present study, we have for the first time shown that diclofenac is a hP2X3R and hP2X2/3R antagonist with micromolar potency. Our results strongly support a competitive antagonism through which diclofenac, by interacting with residues of the ATP-binding site, left flipper, and dorsal fin domains inhibits gating of P2X3R by conformational fixation of the left flipper and dorsal fin domains. In addition, a weaker inhibition of hP2X1R, hP2X4R and hP2X7R was shown. A less potent inhibition of hP2X3R was observed for all other investigated NSAIDs. FFA proved to significantly inhibit hP2X3R, rP2X3R and hP2X7R, questioning its use as a nonselective ion channel blocker, when P2X-mediated currents are under study.

## 2 Materials & Methods

### 2.1 Chemicals

The investigated NSAIDs and most standard chemicals were purchased from Sigma-Aldrich/Merck (Taufkirchen, Germany), if not otherwise specified. ATP sodium salt and α,β-meATP were purchased from Roche Diagnostics (Mannheim, Germany) and Tocris Bioscience (Bristol, United Kingdom), respectively. Collagenase type 2 was purchased from Worthington Biochemical Corp. (Lakewood, USA and distributed by CellSystems, Troisdorf, Germany).

### 2.2 Expression of P2X receptors in *Xenopus laevis* oocytes

Oocyte expression plasmids encoding the wild-type (wt) and His-tagged hP2X2R, hP2X3R, hP2X4R and hP2X7R, the mutant His-^20^RMVL^23^KVIV^23^S^26^N-hP2X1R, S^15^V-rP2X3R and His-S^15^V-hP2X3R were available from previous studies (Hausmann et al., 2006, Hausmann et al., 2014, Obrecht et al., 2019, Wolf et al., 2011). Capped cRNAs of the different P2XR were already available in the research group or were synthesized as previously described (Schmalzing et al., 1991, Stolz et al., 2015). cRNA was injected into collagenase-defolliculated *Xenopus laevis* oocytes in aliquots of 41nl or 23nl (see Suppl. Tbl. 1 for the amount of cRNA used for expression of the indicated P2XR) using a Nanoliter 2000 injector (World Precision Instruments, Sarasota, United States of America) as described previously (Hausmann et al., 2014, Obrecht et al., 2019, Stolz et al., 2015). To express the heteromeric hP2X2/3 receptor, cRNAs encoding His-hP2X2R and wt-hP2X3R were coinjected at a ratio (w/w) of 1:6. Oocytes were stored at 19°C in oocyte ringer solution (ORi^+^) containing 90 mM NaCl, 1 mM KCl, 1 mM CaCl_2_, 1 mM MgCl_2_ and 10 mM HEPES (Carl Roth, Karlsruhe, Germany) adjusted to pH 7.4 with NaOH and supplemented with 50 µg/ml gentamicin (AppliChem, Darmstadt, Germany). The procedures followed for maintaining and surgical treatment of *X. laevis* adults were approved by the governmental animal care and use committee of the State Agency for Nature, Environment and Consumer Protection (LANUV, Recklinghausen, Germany; reference no. 81-02.04.2019.A355), in compliance with Directive 2010/63/EU of the European Parliament and of the Council on the protection of animals used for scientific purposes.

### 2.3 Two-electrode voltage clamp electrophysiology

Ion currents mediated by P2X receptors were evoked by the indicated concentration of ATP or α,β-meATP and were recorded one or two days after cRNA injection at ambient temperature at a holding potential of -60mV by two-electrode voltage clamp (TEVC) as previously described (Hausmann et al., 2006). Calcium-free ORi^-^ solution (90 mM NaCl, 1 mM KCl, 2 mM MgCl_2_, 10 mM HEPES, pH 7.4) was used to avoid bias due to Calcium activated Chloride Channels (CaCC) endogenously expressed in *X. laevis* oocytes (Methfessel et al., 1986, Miledi, 1982). For recordings of the wt-hP2X7R the composition of the ORi^-^ solution was modified according to protocols described previously and contained: 100 mM NaCl, 2.5 mM KCl, 1 mM MgCl_2_, 5 mM HEPES, pH 7.4 (Klapperstück et al., 2000). The oocytes were continuously superfused with ORi^-^ by gravity flow (5-10ml/min). The agonists ATP or α,β-meATP and the investigated NSAIDs were diluted in ORi^-^ on the day of the recording. The following agonist concentrations were used for the different P2X subtypes: 1 μM ATP (hP2X1R mutant, hP2X3R, rP2X3R, S^15^V-hP2X3R, S^15^V-rP2X3R), 10 μM ATP (hP2X2R, hP2X4R), 300 μM free ATP^4-^ (hP2X7R), 1 μM α,β-meATP (hP2X2/3R). A peak current protocol was used to analyse fast- and intermediate desensitizing P2XR subtypes (P2X1R, P2X3R, S^15^V-P2X3R) and a steady-state protocol was used for slowly- or partially-desensitizing P2X subtypes (P2X2R, P2X2/3R, P2X4R). For recordings of the P2X7R a modified steady-state protocol was used (Hausmann et al., 2006). The application of the different bath solutions was controlled by computer-operated magnetic valves controlled by the CellWorks E 5.5.1 software (npi electronic, Tamm, Germany).

### 2.4 Pig Dorsal Root Ganglia Preparation

Dorsal Root Ganglia (DRG) of pigs were sampled according to the 3R criteria for reductions in animal use, as leftovers from previous independent animal studies (e.g. LANUV reference no. 81-02.04.2018.A051). For this purpose, pigs of the German Landrace breed, with an average age of 15 weeks (14.6 SD2.7) and weight of 47.3kg (SD11.2) were euthanized using an overdose of pentobarbital 60 mg/kg body weight. Subsequently, the DRG were collected. The DRG of pigs were transferred on ice and fine excision was performed in ice-cold DMEM F12 medium containing 10 % FBS. DRG were treated with 1mg/ml collagenase P, 1 mg/ml trypsin T1426 and 0,1 mg/ml DNAse for digestion. DRG were cut into small pieces inside the digestion medium for surface enlargement. DRG were incubated in 37 °C, 5 % CO_2_ for 120 minutes ± 30 minutes. Approximately after 60 minutes in digestion medium, DRG were triturated using a glass pipette. After the full incubation time, DRG were triturated three times using glass pipettes with decreasing tip diameter. For further purification, DRG were centrifuged at 500 G and 4 °C twice for four minutes each and the pellets were suspended in DMEM F12 with 10 % FBS. DRG were subsequently separated from the lighter cell fragments and myelin by centrifugation of a Percoll gradient containing a 60 % Percoll and a 25 % Percoll gradient for 20 minutes at 500G. DRG neurons were plated on coverslips coated with poly-D-lysine (100 μg/ml), laminin (10 μg/ml) and fibronectin (10 μg/ml). Neurons were then cultured in neurobasal A medium supplemented with B27, penicillin, streptomycin and L-glutamine and used for voltage-clamp recordings after 12-72 hours in culture.

### 2.5 Whole-Cell Patch-Clamp Recordings of Pig DRG Neurons

Whole-cell voltage clamp recordings of DRG neurons were performed using glass electrodes with micropipette tip resistances of 1,3-3,5 MΩ, pulled and fire-polished with a Zeitz DMZ-puller. The intracellular solution contained 10 mM NaCl, 140 mM CsF, 10 mM HEPES, 1 mM EGTA, 5 mM glucose, 5 mM TEA-Cl (adjusted to pH 7.3 using CsOH). The extracellular bathing solution contained 140 mM NaCl, 3 mM KCl, 1 mM MgCl_2_, 1mM CaCl_2_, 10 mM HEPES and 20 mM glucose (adjusted to pH 7.4 using NaOH). The liquid junction potential was corrected by -7.8 mV. Membrane currents were measured at room temperature with a holding potential of -77.8 mV using a HEKA EPC-10 USB amplifier. 10 μM α,β-methylene ATP and 100 μM Diclofenac were applied using a gravity-driven perfusion system during the recordings. PatchMaster/FitMaster software (HEKA Electronics) and IgorPro (WaveMetrics) were used for data acquisition and analysis. Signals were digitized at a sampling rate of 5 kHz. The low-pass filter frequency was set to 10 kHz. Series resistance compensation was between 2.5 and 11.1 MΩ.

### 2.6 Data Analysis

The recorded TEVC-currents were analyzed using CellWorks Reader 6.2.2 (npi electronic, Tamm, Germany) and Microsoft Excel (Microsoft Corporation, Redmond, USA). The displayed current traces were generated with Igor Pro 6.21 (WaveMetrics, Portland, USA) and edited with Microsoft PowerPoint (Microsoft Corporation, Redmond, USA). To generate concentration-response curves, non-linear regression analysis was performed using GraphPad Prism 5 (GraphPad Software, San Diego, United States of America).

Antagonist concentration–response data and IC_50_ values were calculated by normalizing ATP-induced responses to the control responses (recorded in the presence and absence of antagonist, respectively). The four-parameter Hill equation (Eq. (1)) was iteratively fitted to data collected from a minimum of four independent repeat experiments to obtain antagonist concentration– response curves and IC_50_ values.

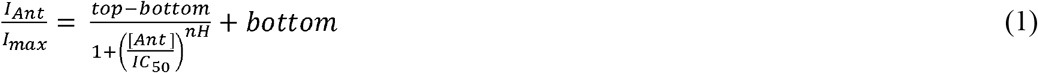

I_max_ is the control response in the absence of antagonist, I_Ant_ is the response at the given antagonist concentration [Ant], and IC_50_ is the antagonist concentration that causes 50% inhibition of the response elicited by a given agonist concentration. The ratio between the response in presence of a certain antagonist concentration and the control response in absence of the antagonist is indicated as “% control current”.

In case of fast-desensitizing P2XR subtypes (P2X1R, P2X3R), ATP is applied five times in repetition and the ATP-induced current amplitude in the presence (after pre-incubation) of the antagonist (fourth application) is compared to the arithmetic mean of flanking control ATP-induced current amplitudes in the absence of the antagonist (third and fifth ATP application). Since some of the investigated NSAIDs showed an enduring, potentially irreversible inhibitory effect on the current amplitude, only the preceding (third) ATP-induced current amplitude was used to calculate the control current. The typical run-down of current amplitudes between consecutive, repetitive ATP applications was considered by applying a correction factor. The correction factor was calculated as the ratio of the fourth ATP-induced current amplitude to the third amplitude (Eq. (3)), when the experiment was performed in absence of the antagonist. When the experiment was performed in presence of the antagonist, the ATP-induced current amplitude of the preceding current (third amplitude) was multiplied by this correction factor / quotient (Eq. (4)) to obtain a control current corrected for the run-down effect. Since the magnitude of the run down varies from day to day and batch to batch of oocytes, the correction factor was determined on each day of experiments and was calculated as the arithmetic mean of several recordings on each day.

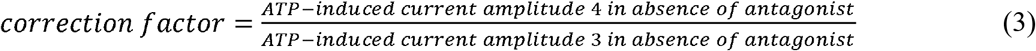

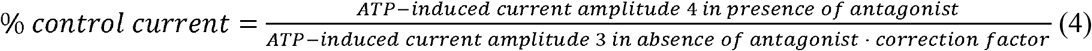

In case of the hP2X7R, the control current in absence of the antagonist had to be extrapolated (Suppl. Fig. 4, 5) presuming a linear increase of permeability during continuous ATP application (North, 2002), since the permeability of the receptor is affected by the antagonist.

The IC_50_ values are displayed as geometric means with 95% confidence intervals (95% CI). All other values (including % control current or % inhibition) are presented as arithmetic means ± SEM.

### 2.7 hP2X3R X□ray structure□based molecular dynamics simulations and evaluation of diclofenac binding

The human ionotropic cation-selective ATP receptor P2X3 was modeled based on its structure in the open state (PDBID: 5SVK) and apo-closed state (PDBID: 5SVJ) (Mansoor et al., 2016) and embedded in a 1-palmitoyl-2-oleoyl-phosphatidylcholine (POPC) bilayer using the g_membed functionality (Wolf et al., 2010) in GROMACS. Modeller was used to build the single-point mutant L191A of the P2X3 apo-closed state in order to make the supposed binding cavity of diclofenac easier accessible to be able to simulate the binding event within the µs-time-scale of Molecular Dynamics simulations (Fiser and Sali, 2003). The system was simulated in GROMACS (Abraham et al., 2015) version 2021 using a time step of 2 fs.

A pressure of 1 bar was applied semi-isotropically with a Berendsen barostat (Berendsen et al., 1984) using a time constant of 5 ps. A temperature of 310 K was maintained with a velocity-rescaling thermostat (Bussi et al., 2007). Van der Waals interactions were calculated with the Lennard-Jones potential and a cutoff radius of 1.2 nm, with forces smoothly switched to zero in the range of 1.0−1.2 nm and no dispersion correction. The protein was described by the CHARMM36m (Huang et al., 2017) force field, lipids by the CHARMM36 force field (Klauda et al., 2010), and water by the TIP3P model (Jorgensen et al., 1983).

Na^+^ and Cl^−^ were added to give a bulk concentration of approximately 50 mM NaCl. Three diclofenac molecules were added per system and fitted onto AF-219 of aligned 5SVQ. 7 independent systems each of the wildtype P2X3 apo-closed state and L191A apo-closed state were simulated for more than 200 ns each, and were preceded by equilibration for about 200 ns: first with restraints on all heavy atoms and lipids in the z-direction, second on all heavy atoms, and third on backbone atoms only. All trajectories that showed a stable binding of diclofenac were selected and clustered with GROMACS tool gmx clustern and gromos algorithm. Cut-off for RMSD differences in a cluster was set to 0.3 nm.

Initial force-field parameters for diclofenac were generated according to the CHARMM generalized force-field (CGenFF) (Vanommeslaeghe et al., 2010, Vanommeslaeghe and MacKerell, 2012, Vanommeslaeghe et al., 2012, Yu et al., 2012), using the CHARMM-GUI webserver (https://charmm-gui.org/). The initial molecular geometry and charge assignments were further optimized with the force-field toolkit (ffTK) (Mayne et al., 2013) version 2.1 plugin for the visual molecular dynamics (VMD) version 1.9.4a57 analysis suite (Humphrey et al., 1996). The ffTK program provides a workflow of quantum-mechanical calculations using ORCA (Neese et al., 2020) 5.0.3, followed by Newtonian optimizations using the nanoscale molecular dynamics (NAMD) (Phillips et al., 2020) engine. An initial parameter file is generated in ffTK by analogy using the protein structure file (psf) and protein coordinate file generated by CHARMM-GUI. The initial molecular geometry was optimized with ORCA at the MP2/6-31G* level of theory. After the geometry-optimization had converged, atomic partial charges were approximated with ORCA by calculating water-interaction energies at the HF/6-31G* level of theory. Aliphatic and aromatic hydrogens were assigned partial charges of 0.09 and 0.115 respectively – only hydroxyl hydrogens were optimized. To account for the positive charge associated with the dipole created by halogens known as alpha-holes, a lone-pair particle (LP) was added automatically via CHARMM-GUI (Pang et al., 2020). New bonded parameters for diclofenac only contained two dihedral terms that, which are consistent with the CHARMM36-ForceField, were used for diclofenac simulations – dihedral bonds were not further optimized due to accordance with having a bond energy-penalty of less than 50 (unitless penalties as provided by the CGenFF program). Detailed instructions for using the most updated ffTK with support for the open-source quantum chemistry package, ORCA, can be found at the ffTK website (https://www.ks.uiuc.edu/Research/vmd/plugins/fftk), and the updated tutorial (https://www.ks.uiuc.edu/~mariano/fftk-tutorial.pdf).

## 3 Results

### 3.1 Validation of the inhibitory effect of various NSAIDs on P2X3R-mediated currents

In a previous screening of 2000 approved drugs, natural products and bioactive substances, various NSAIDs were found to inhibit S15V-rP2X3R-mediated currents (Obrecht et al., 2019). These included diclofenac, flunixin meglumine, meclofenamic acid and niflumic acid, where diclofenac showed the greatest inhibitory effect (> 80 % inhibition) of the screened NSAIDs (Obrecht et al., 2019). To validate the screening results, we characterised the potency of diclofenac, flunixin and meclofenamic acid using TEVC on *Xenopus laevis* oocytes heterologously expressing S^15^V-rP2X3R and His-S^15^V-hP2X3R. Instead of niflumic acid, we decided to investigate the structurally related flufenamic acid (FFA) due to its additional use in research as a nonselective ion channel blocker (Guinamard et al., 2013). Furthermore, we included the NSAIDs ibuprofen and naproxen in our investigations, because these are used extensively in daily practice. The structural formulas of the NSAIDs investigated are shown in Figure 1 (Fig. 1, compounds 1-6).

**Figure 1.**
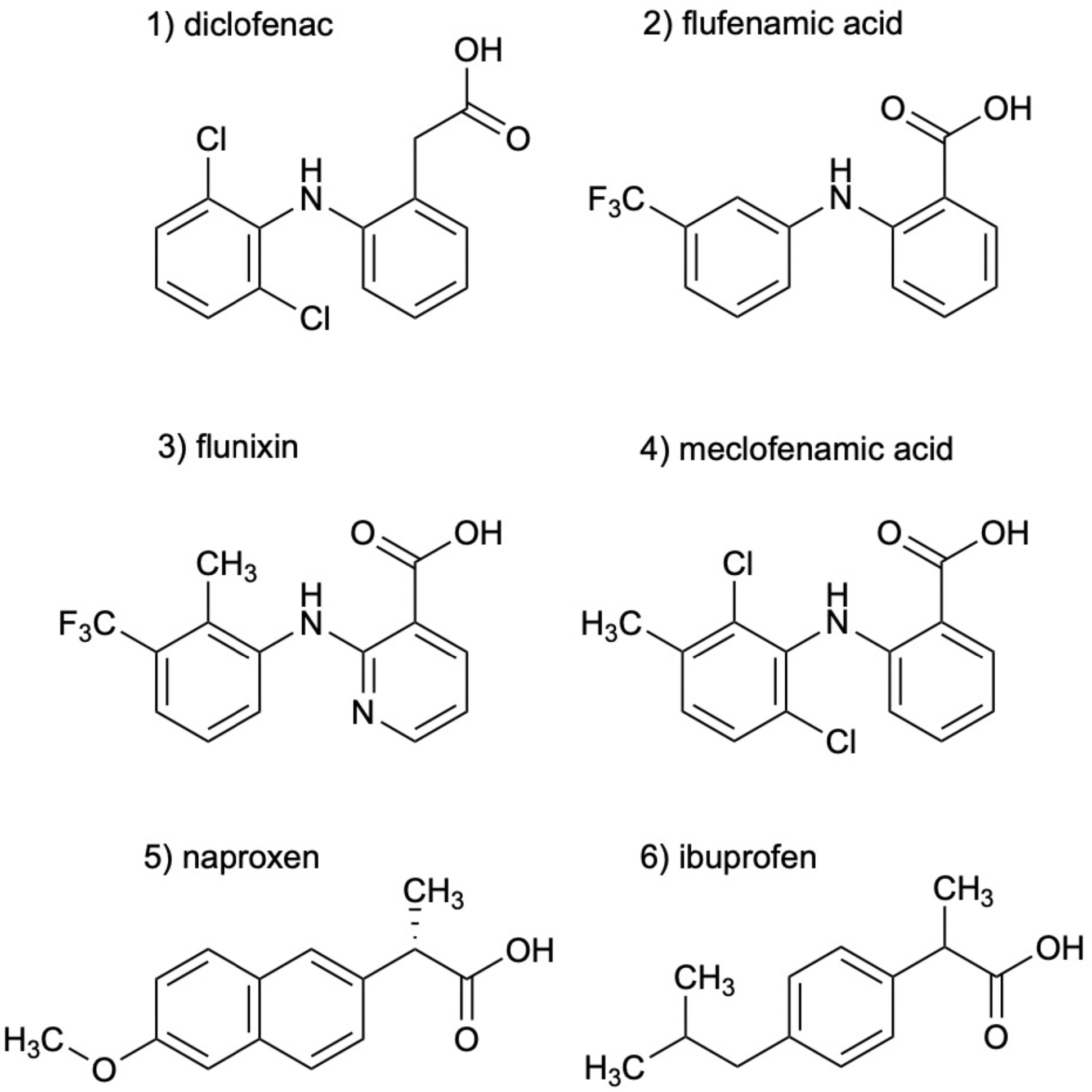
Chemical structures of the investigated NSAIDs (1-6).

A peak current protocol including a 30 s preincubation of the antagonist was applied (Suppl. Fig. 1) to determine the inhibitory effect of the NSAIDs at the P2X3Rs. The current amplitude in presence of the antagonist was compared to the preceding control current in absence of the antagonist. The inhibitory effect of diclofenac, FFA and flunixin was not reversible by the following washout, which was reflected in a reduced amplitude of the subsequent control current that could not be explained by the run down alone (c.f. Fig. 2 A). Due to this potentially irreversible inhibitory effect, the subsequent amplitude in absence of the antagonist did not provide a suitable reference and was not taken into account to calculate the control current.

**Figure 2.**
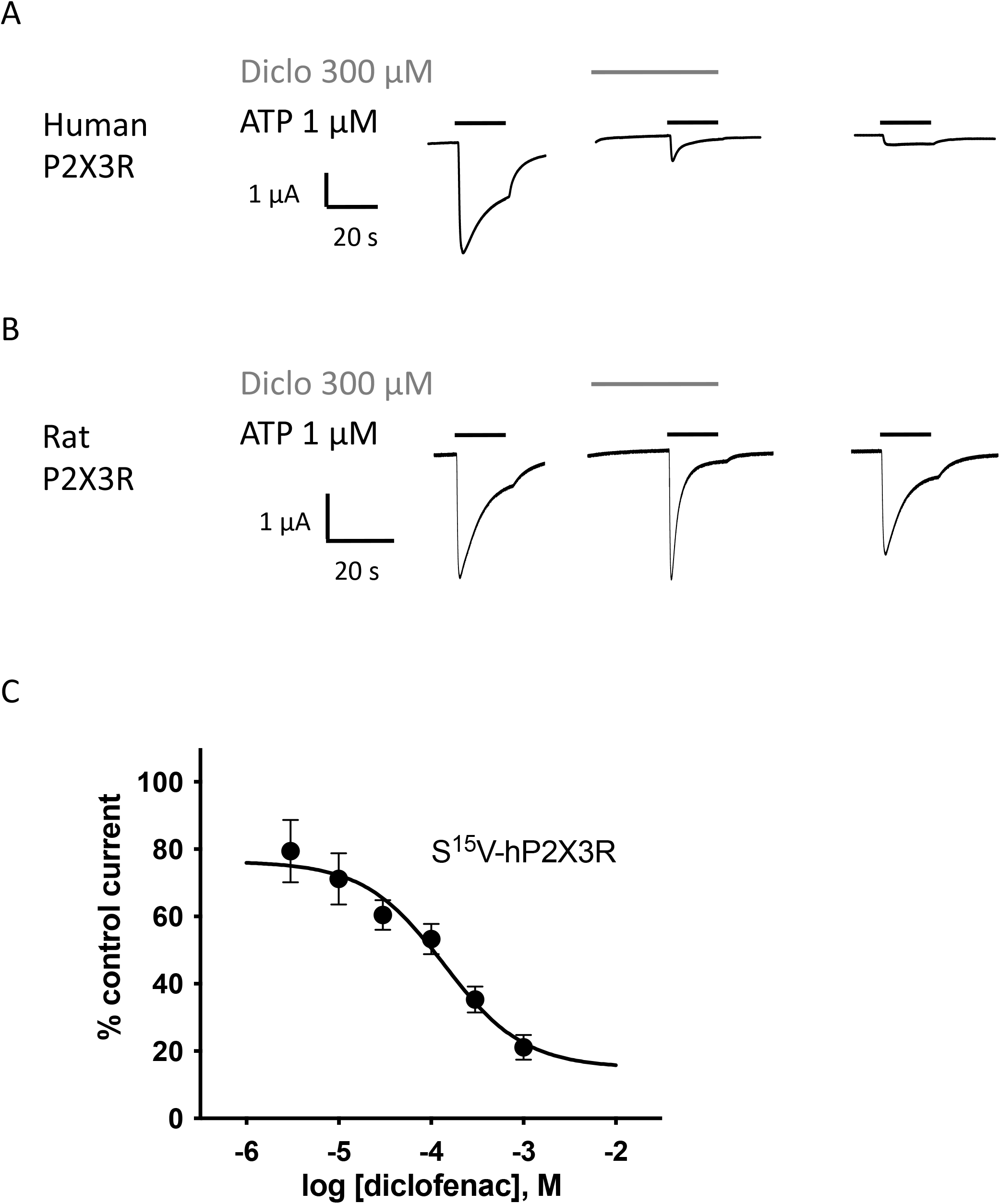
Diclofenac inhibition of ATP-induced currents of the P2X3R is concentration dependent. The effect of diclofenac on ATP-induced (1 μM) currents through S15V-hP2X3R (A) and S15V-rP2X3R (B) expressed in *X. laevis* oocytes was analyzed by TEVC electrophysiology. (A, B) Representative original current traces show the effects of 300 µM diclofenac on ATP-induced (black bars) currents of the indicated P2X3R. To ensure a binding equilibrium was reached, diclofenac was pre-incubated for 30 s before adding ATP (grey bar). (C) The resulting concentration–inhibition curve of diclofenac at the hP2X3R exhibited an IC_50_ of 138.2 µM (95% CI: 46.0 - 415.1 µM). Data points represent the means and SEM.

All investigated NSAIDs were less effective at the rat P2X3R than at the human P2X3R (Fig. 2 A, B) or even had no effect at all on rat P2X3R, thus we decided to focus our further investigations on human P2X receptors. Concentration-response analysis revealed that ATP-evoked hP2X3R-mediated responses were inhibited by various NSAIDs. Diclofenac proved to be the most effective antagonist with an IC_50_ value of 138.2 µM (95% CI: 46.0 - 415.1 µM; Fig. 2 C) and a maximum inhibition of ∼ 80% at a concentration of 1mM (Tbl. 1). FFA proved to inhibit hP2X3R-mediated currents with a lower potency (IC_50_ value of 208.8 µM; 95% CI: 94.8 - 459.9 µM; Tbl. 1) in comparison to diclofenac. Flunixin, which is mainly used in veterinary medicine, had a greater potency for the hP2X3 receptor than FFA and diclofenac (IC_50_ value of 32.4 µM; 95% CI: 11.6 - 90.2 µM; Tbl. 1), but a maximum inhibition of only 53% at a concentration of 1mM was observed, indicating a lower efficacy of flunixin. By contrast, only a weak inhibition of hP2X3R was observed for ibuprofen, meclofenamic acid and naproxen. The current amplitude in presence of 100 μM meclofenamic acid, naproxen or ibuprofen was reduced by a maximum of 15-18% suggesting an estimated IC_50_ value of > 300 µM (Tbl. 1). Due to their low inhibitory potency (ibuprofen, meclofenamic acid and naproxen) or low efficacy (flunixin) (c.f. Tbl. 1) these were not investigated further. Thus, only diclofenac - being the most effective antagonist - and FFA due to its additional use in research as a nonselective ion channel blocker (Guinamard et al., 2013) were further analyzed.

**Table 1:** Concentration-response analysis of NSAIDs at the S15V-hP2X3R expressed in *X. laevis* oocytes. n.d., not determined.

|  | IC<sub>50</sub>, μM | 95 % CI IC<sub>50</sub> | Max. Inhibition at 1 mM | n |
| --- | --- | --- | --- | --- |
| Diclofenac | 138.2 | 46.0 – 415.1 | 78.9 ± 3.7 % | 10 |
| Flufenamic acid | 221.7 | 98.9 – 496.8 | 69.8 ± 3.8 % | 10 |
| Flunixin | 32.4 | 11.6 – 90.2 | 53.7 ± 4.8 % | 8 |
| Ibuprofen | > 300 | n.d. | n.d. | 8 |
| Meclofenamic acid | > 300 | n.d. | n.d. | 9 |
| Naproxen | > 300 | n.d. | n.d. | 9 |

### 3.2 Characterisation of the potency and selectivity of diclofenac at P2X receptors by TEVC

Selectivity profiling of diclofenac was performed at the hP2X1R, hP2X2/3R, hP2X2R, hP2X4R and hP2X7R (Fig. 3 B). To analyze the heteromeric hP2X2/3R the ATP derivate α,β-meATP was used (Fig. 3 A) to evoke hP2X2/3R-mediated currents, because it does not activate the homotrimeric hP2X2R. Currents mediated by the homotrimeric hP2X3R can be neglected due to its strong desensitization and run down, when α,β-meATP is applied repetitively in short intervals (Bianchi et al., 1999, North, 2002). The desensitization kinetics of the heterotrimeric hP2X2/3 receptor resemble those of the homotrimeric hP2X2 receptor, therefore the steady-state protocol was used (Fig. 3 A) (North, 2002). Diclofenac antagonized the heteromeric P2X2/3R with the highest potency of 76.7 µM (95% CI: 64.6 - 91.2 µM) and showed the following rank order of its potencies at P2XR subtypes: hP2X2/3R > hP2X3R > hP2X1R > hP2X4R. All IC_50_ values are summarized in table 2 (Tbl. 2). By contrast, at the hP2X2R diclofenac did not antagonize ATP-evoked P2X2R-mediated responses, but presence of 1 mM of diclofenac exhibited a 1.7-fold increase of the P2X2R responses and thus showed a potentiating effect at the P2X2R (Suppl. Fig. 3 A).

**Figure 3.**
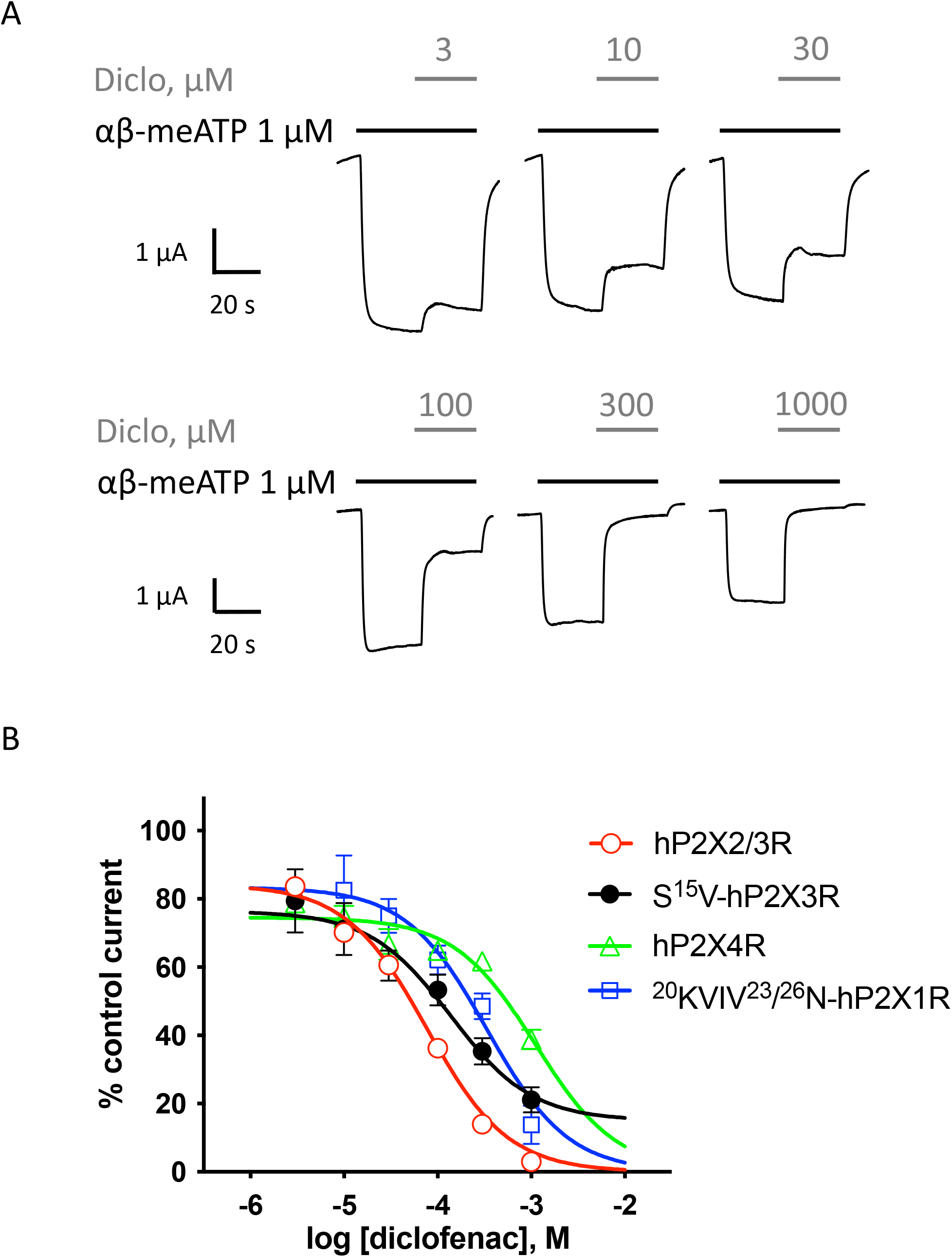
Effect of diclofenac at heteromeric hP2X2/3R and selectivity profiling. (A) Representative original current traces show the effects of 3 - 1000 µM diclofenac (grey bars) on α,β-meATP-induced (1 μM, black bars) currents through heteromeric hP2X2/3R expressed in *X. laevis* oocytes. (B) Concentration–inhibition curve of diclofenac at the indicated P2XR isoforms are shown as obtained from TEVC measurements using a peak current (P2X1R, P2X3R) or steady state (P2X2/3R, P2X4R) protocol (c.f. Suppl. Figs. 1/2). All IC_50_ values are summarized in Table 2.

**Table 2:**
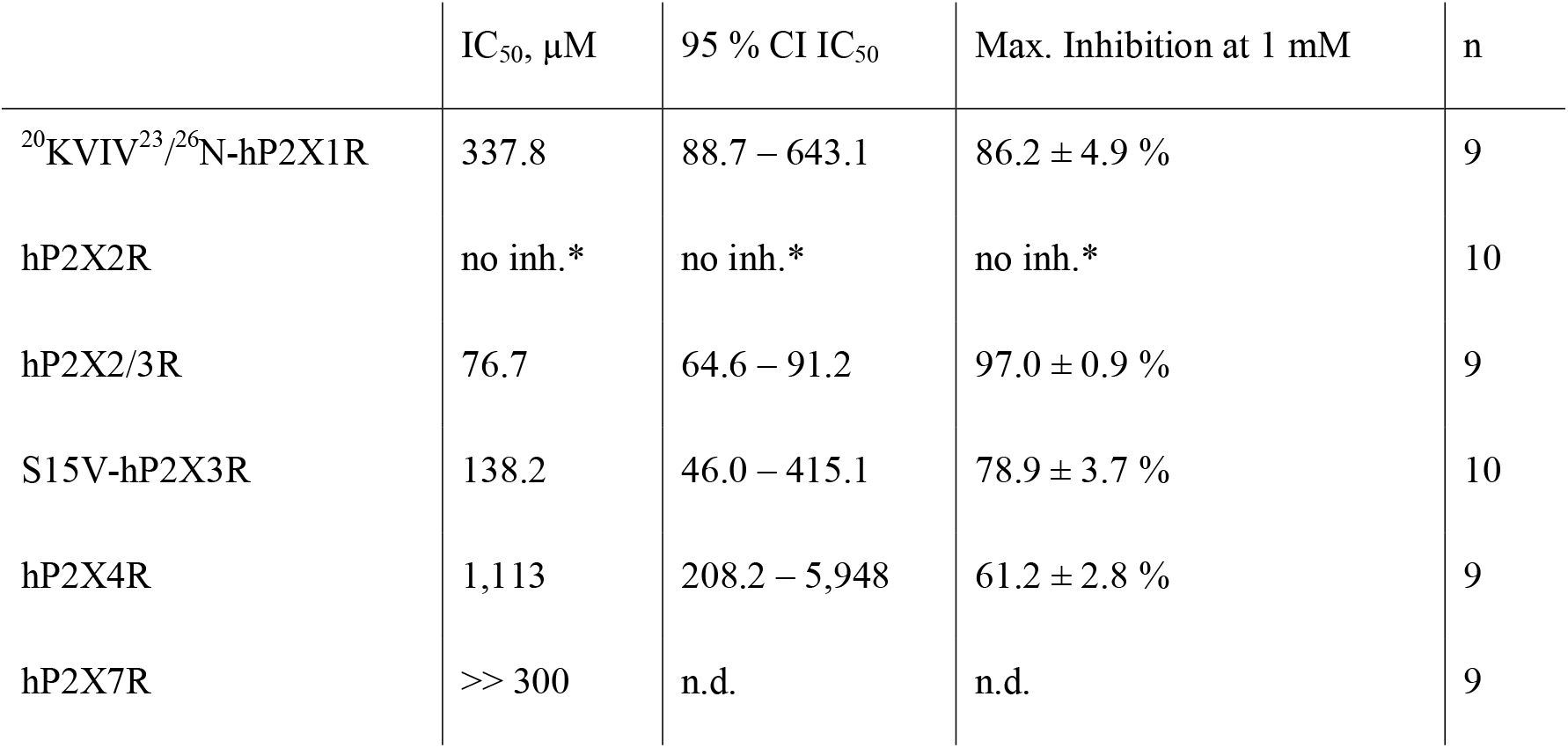
Concentration-response analysis of diclofenac at the indicated hP2XR subtypes expressed in *X. laevis* oocytes. *, no inhibition, but potentiation (c.f. Suppl. Fig. 3); n.d., not determined.

|  | IC<sub>50</sub>, μM | 95 % CI IC<sub>50</sub> | Max. Inhibition at 1 mM | n |
| --- | --- | --- | --- | --- |
| <sup>20</sup>KVIV<sup>23</sup>/<sup>26</sup>N-hP2X1R | 337.8 | 88.7 – 643.1 | 86.2 ± 4.9 % | 9 |
| hP2X2R | no inh.* | no inh.* | no inh.* | 10 |
| hP2X2/3R | 76.7 | 64.6 – 91.2 | 97.0 ± 0.9 % | 9 |
| S15V-hP2X3R | 138.2 | 46.0 – 415.1 | 78.9 ± 3.7 % | 10 |
| hP2X4R | 1,113 | 208.2 – 5,948 | 61.2 ± 2.8 % | 9 |
| hP2X7R | >> 300 | n.d. | n.d. | 9 |

In summary, diclofenac was shown to be more potent at hP2X2/3R (IC_50_ 76.7 µM; 95% CI: 64.6 - 91.2 µM) than at hP2X3R (IC_50_ 138.2 µM; 95% CI: 46.0 - 415.1 µM). However, it should be noted that the use of the different agonists (α,β-meATP hP2X2/3R; ATP hP2X3R) complicates the assessment of a quantitative comparative analysis.

To assess the effect of diclofenac on the non-desensitizing hP2X7R a modified steady-state protocol was used. For recordings of the P2X7R, most scientists use divalent free solutions such as ORi-supplemented with 100 µM flufenamic acid (FFA) as an unselective ion channel blocker to inhibit nonspecific chloride conductance in the absence of divalent ions (Hülsmann et al., 2003, Weber et al., 1995). However, since FFA is one of the investigated NSAIDs, this supplement did not seem to be reasonable. Therefore, the composition of the ORi-solution was modified as follows: 100 mM NaCl, 2.5 mM KCl, 1 mM MgCl_2_, 5 mM HEPES, pH 7.4 and according to former protocols (Klapperstück et al., 2000) an free ATP^4-^concentraion of 300 µM was adjusted. 300µM of diclofenac reduced the ATP^4-^-induced current amplitude by approximately 33% (Suppl. Fig. 4), which suggested an estimated IC_50_ value of > 300 µM for diclofenac at the hP2X7R. Since such high concentrations of diclofenac are clinically irrelevant, we refrained from performing a concentration-response analysis.

### 3.3 Mechanism of action of diclofenac

Although the S^15^V mutant of P2X3R desensitizes slowly (Hausmann et al., 2014), the desensitization may still prevent reliable assessment of the mechanism of antagonism (Hausmann et al., 2014). Thus, the non-desensitizing heteromeric hP2X2/3R was used assess the mechanism of action of diclofenac, which was also inhibited by diclofenac with the highest potency. To this end, the extent of inhibition of the heteromeric hP2X2/3R by 30 µM diclofenac was determined using α,β-meATP concentrations of 1 µM or 30 µM as an agonist. We refrained from determining full agonist concentration-response curves at the hP2X2/3R, because simultaneous activation of homomeric hP2X2R occurs when hP2X2 and hP2X3 subunits are co-expressed and agonist concentration exceeds 30 µM α,β-meATP. 30 µM diclofenac inhibited the 1 µM or 30 µM α,β-meATP-induced current responses of the heteromeric hP2X2/3R by 44.8 ± 21.9 % or 25.1 ± 10.1 %, respectively (Fig. 4 A). Thus, inhibition by 30 µM diclofenac could be overcome by increasing concentrations of the agonist α,β-meATP, indicating competition of diclofenac and α,β-meATP at the hP2X2/3R.

**Figure 4.**
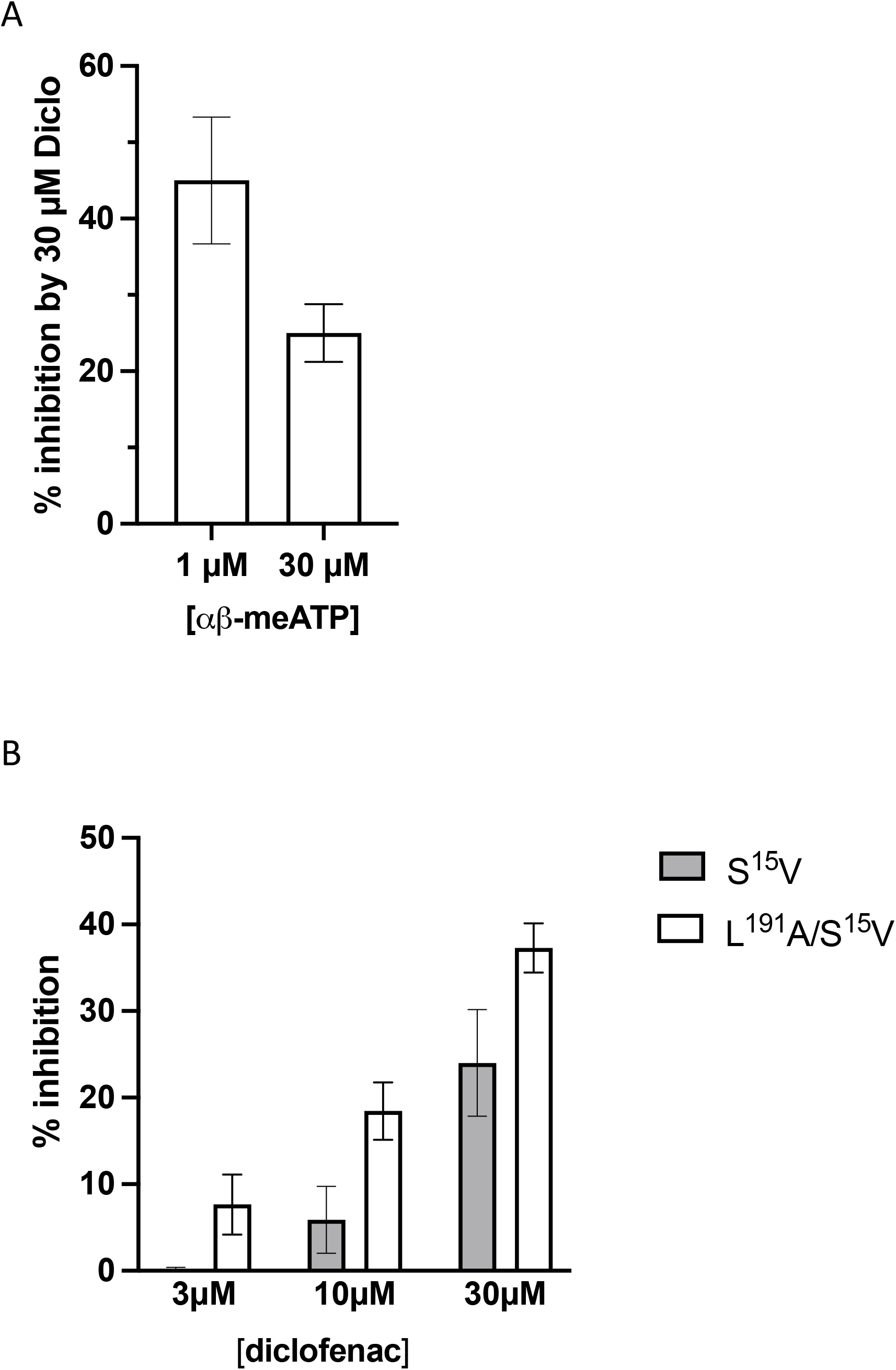
Diclofenac inhibition is modulated by the agonist concentration and the L^191^A mutant. (A) Bar graph showing the inhibition of the heteromeric hP2X2/3R by 30 µM diclofenac as determined with agonist concentrations of 1 µM or 30 µM α,β-meATP as indicated (1 µM α,β-meATP, 44.8 ± 21.9 %, n = 7; 30 µM α,β-meATP, 25.1 ± 10.1 %, n = 7). Inhibition by 30 µM diclofenac could be overcome by increasing concentrations of α,β-meATP. (B) Comparative analysis of diclofenac inhibition of 10 µM ATP-induced currents of the S^15^V-hP2X3R and L^191^A/S^15^V-hP2X3R. Bar graphs showing the inhibition of the indicated receptor by 3, 10 or 30 µM diclofenac (% inhibition ± SEM: 3 µM diclofenac: S^15^V -1.6 ± 2.02 %, n = 8 and L^191^A/S^15^V 7.7 ± 3.5 %, n = 17; 10 µM diclofenac: S^15^V 5.9 ± 3.9 %, n = 12 and L^191^A/S^15^V 18.4 ± 3.3 %, n = 16; 30 µM diclofenac: S^15^V 24.0 ± 6.2 %, n = 8 and L^191^A/S^15^V 37.3 ± 2.8 %, n = 20. The L^191^A mutant is inhibited to a greater extent by diclofenac.

To further support the competitive nature of the inhibition and to exclude the possibility that diclofenac does bind in the negative allosteric site of hP2X3R as do other modulators (or negative allosteric antagonists) such as gefapixant (formerly AF-219) (Wang et al., 2018) or ATA (Obrecht et al., 2019), we examined mutations of amino acid residues in the alloSite with respect to the effect of diclofenac. We have analyzed the L^191^F, N^190^A and the G^189^R mutants (in the background of the S^15^V-hP2X3R (Obrecht et al., 2019)) of the negative allosteric binding site described in previous studies (Obrecht et al., 2019, Wang et al., 2018). These were inhibited by 100 µM diclofenac to a similar extent than the S^15^V-hP2X3R, suggesting that the negative allosteric site of hP2X3R is not the binding site of diclofenac. Furthermore, in contrast to the findings for negative allosteric antagonists gefapixant/AF-219 (Wang et al., 2018) or ATA (Obrecht et al., 2019) the L^191^A/S^15^V-hP2X3R mutant was inhibited to a marked greater extent by diclofenac compared to the S^15^V-hP2X3R (Fig. 4 B). The competitive nature of the diclofenac inhibition as well as FFA inhibition of the L^191^A/S^15^V-hP2X3R mutant can be derived from Suppl. Fig. 6, which illustrates representative original current traces of the L^191^A/S^15^V-hP2X3R showing the effects of 10 µM diclofenac or 30 µM FFA on ATP-induced currents: the initial inhibition by diclofenac or FFA at the beginning of the co-application could be overcome by prolonged ATP co-application, indicating competitive binding of ATP and the antagonist to the ATP-binding site.

To investigate the binding mode of diclofenac and to shed light on a possible inhibition mechanism, we performed extensive all-atom molecular dynamics simulations of hP2X3R (Fig. 5 A) embedded in a lipid bilayer and surrounded by a physiological NaCl-based solution. Since the L^191^A mutation appears to facilitate diclofenac binding in our experiments, we initially assumed that diclofenac interacts with the receptor at a similar site like the allosteric inhibitor AF-219 (Wang et al., 2018), although L^191^A was shown to reduce binding of this compound. Therefore, we placed diclofenac molecules at the position of AF-219 (Wang et al., 2018) and investigated how diclofenac reorients in equilibrium simulations over hundreds of nanoseconds.

**Figure 5.**
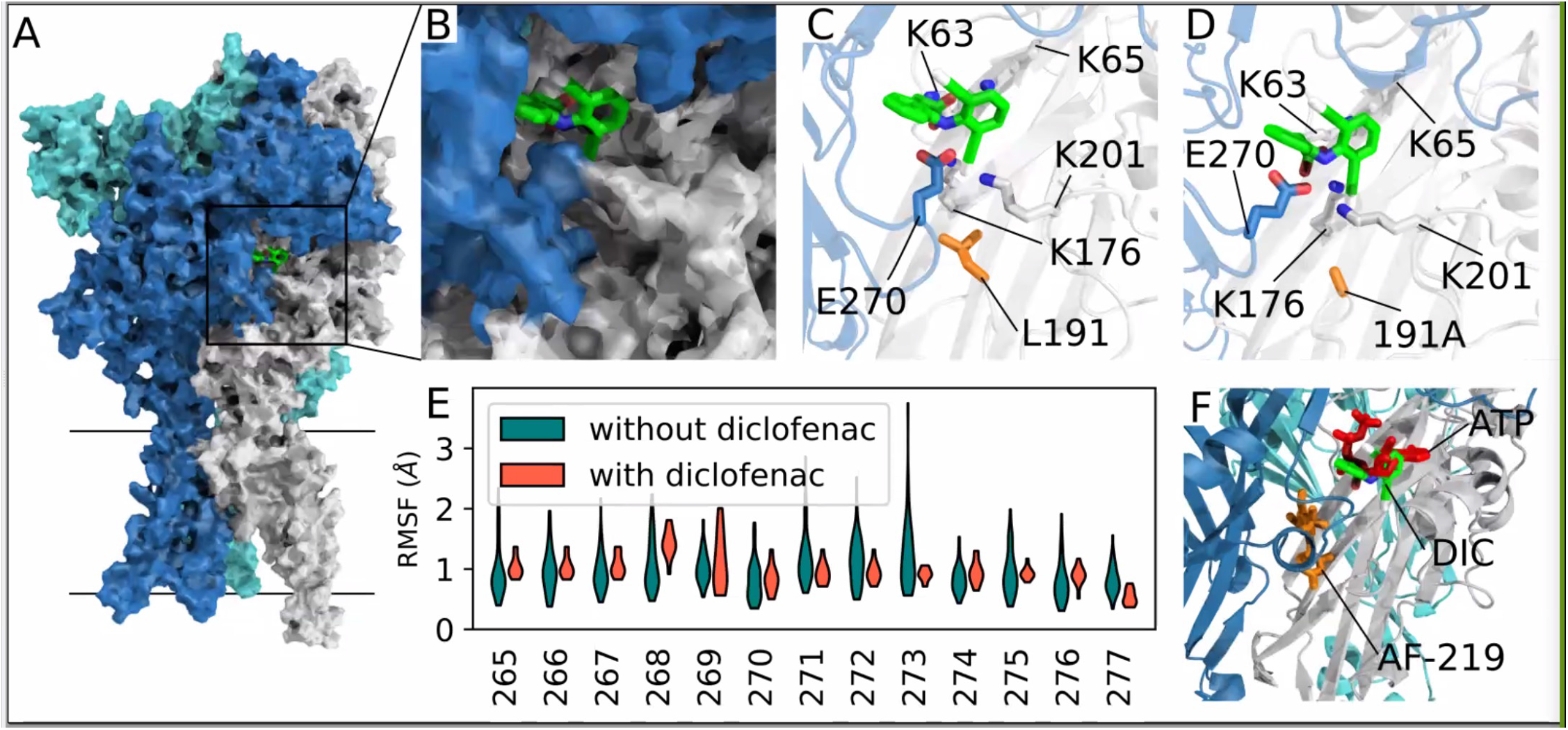
Comepetitive mechanism of action as revealed by extensive all atom molecular dynamics simultations. (A) Surface representation of apo-state P2X3 (PDB: 5SVJ) with bound diclofenac (as obtained in our MD simulations) in the agonist binding pocket between two adjacent subunits. (B) Close-up of the diclofenac-binding pocket as in (A). (C) Average structure of the most frequently observed binding pose of diclofenac in P2X3 wildtype. Interacting residues within 4 Å distance are shown as sticks. (D) Same as D in the apo state L191A mutant. The average structure of the first cluster of the independent simulations of the mutant show a nearly identical binding pose. (E) RMSF (root-mean-square fluctuation) of the left flipper domain for apo state P2X3 wildtype with and without bound diclofenac to the agonist binding pocket. (F) Open state P2X3 with bound ATP (PDB: 5SVK)overlaps with bound diclofenac when being aligned with apo-closed state. AF-219 (PDB: 5YVE) binds to a lower cavity.

In seven independent simulation replicas, we consistently observed diclofenac to alternatingly form salt-bridge interactions between its carboxyl group and residues K^65^, K^63^, and K^176^. Furthermore, diclofenac binding was stabilized by a hydrogen bond with E^270^ and interactions between one of diclofenac’s chlorine atoms at K^176^ and K^201^ (Fig. 5 B, C). A similar diclofenac binding pose was observed in simulations of L^191^A-hP2X3R. We speculate that removal of the bulky hydrophobic sidechain of L^191^ may facilitate diclofenac binding by creating an energetically more favorable environment (Fig 5 D). We then calculated the root-mean-squared fluctuations of the loop of the left flipper domain (residue stretch 265–277, hP2X3 numbering) and observed a reduction in flexibility upon diclofenac binding. Thus, it appears that diclofenac binding rigidifies this region and may thereby impairs allosteric communication between the ATP binding site and the lower body and transmembrane domains (Fig. 5 E). Summarizing, our simulations resolved the binding pose of diclofenac, which is nearby but distinct from the binding pose of the allosteric inhibitor AF-219, and partially overlaps with ATP, suggesting a partially competitive inhibition mechanism (Fig. 5 F).

### 3.4 Effect of diclofenac at native P2XRs in DRG neurons

To examine whether diclofenac is capable of inhibiting native P2X3-subunit containing receptors of nociceptive neurons with similar potency as oocyte-expressed recombinant hP2X2/3Rs and P2X3Rs, DRG neurons of pigs (3 - 4 month old) were analyzed. Currents elicited by 10 μM α,β-meATP were found in medium sized (∼ 35 - 60 μm diameter) porcine DRG neurons. 10 μM α,β-meATP was applied repeatedly every 3 min for 3 s duration onto cultured porcine DRG neurons (Fig. 6 A, B). Whole cell currents elicited by α,β-meATP appeared as a slower activating and non-desensitizing phenotype mediated by heteromeric P2X2/3Rs. These were inhibited by 100 μM diclofenac (pre-equilibrated for 20 s) by 70.5 ± 35.8 % (n = 11) (Fig. 6 C). Thus, diclofenac inhibited native pig P2X2/3Rs expressed in medium sized DRG neurons to a similar extent than hP2X2/3Rs heterologously expressed in *X. laevis* oocytes (c.f. Fig. 3).

**Figure 6.**
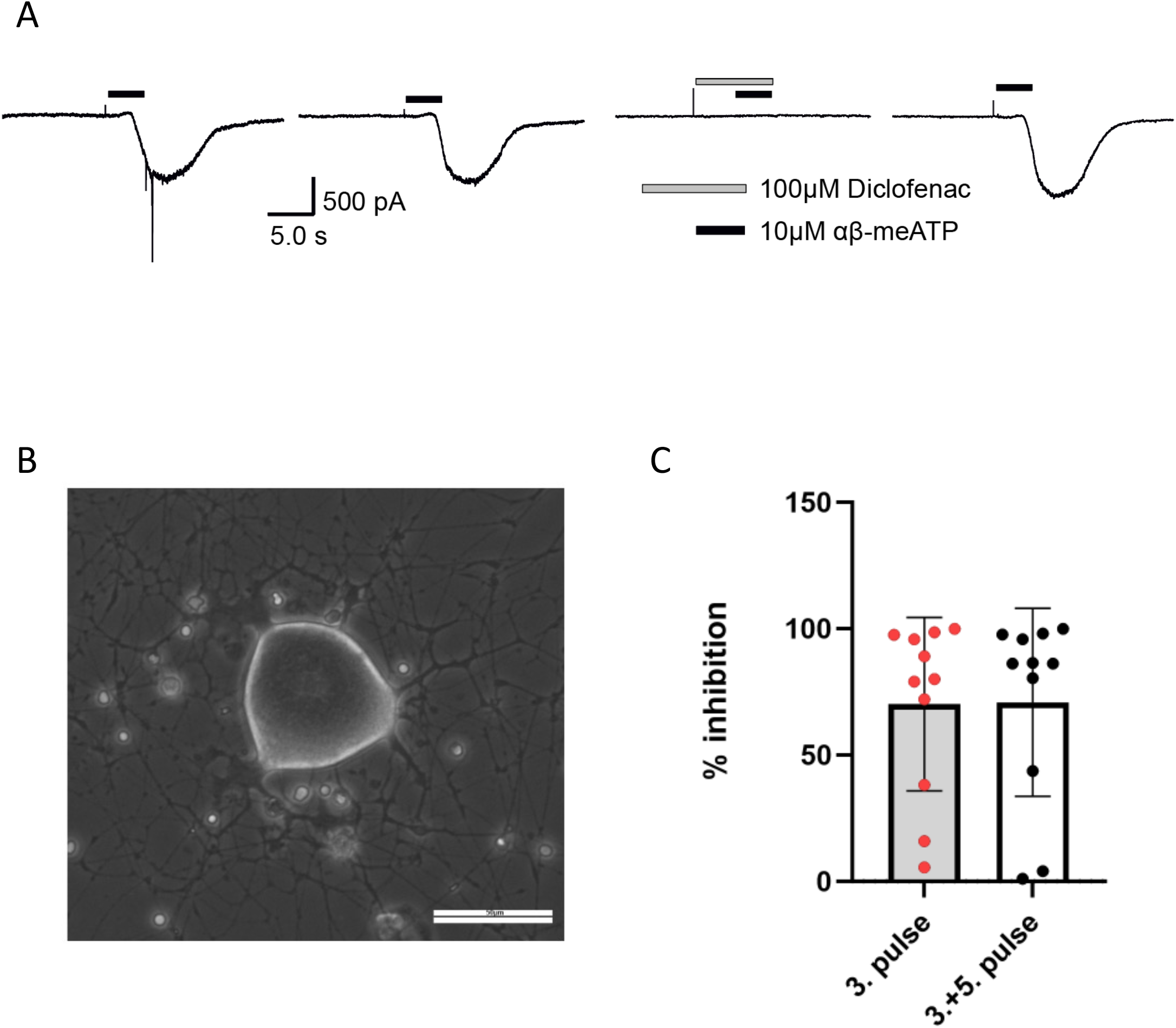
Diclofenac inhibition of P2X2/3R currents in dissociated porcine DRGs. (A) Representative original current traces of one porcine DRG neuron exposed four times for 3s to 10 µM α,β-meATP in 3 minutes intervals. Please note that the neurons were exposed to five applications and that the first application is not shown here. Before the forth application, 100 µM diclofenac was pre-incubated for 20s before 10 µM α,β-meATP and 100 µM diclofenac was co-applied. (B) Representative picture of dissociated porcine DRGs at day 2 in culture. In the center a middle sized DRG neuron is visible. Scale bar = 50 µm. (C) Bar graphs showing the relative diclofenac inhibition as calculated by the quotient of the max. α,β-meATP-induced peak current amplitude of the forth application (in presence of diclofenac) vs. the preceding 3rd ATP application (mean Block = 70.2 ± 34.3 %) (left bar) or vs. the mean of the preceding (3^rd^) and following (5^th^) α,β-meATP application (mean Block = 70.9 ± 37.2 %) (right bar).

### 3.5 Characterisation of the potency and selectivity of FFA at selected P2X receptors

Due to its common use in research as a nonselective ion channel blocker (Guinamard et al., 2013), FFA was further characterized at selected P2XRs.

Concentration-response analysis revealed a concentration-dependent inhibition of hP2X3R- and rP2X3R-mediated currents by micromolar concentrations of FFA. IC_50_ values of 221.7 µM (95% CI: 98.9 - 497 µM) and 264.1 µM (95% CI: 56.9 – 612 µM) were determined at the hP2X3R and rP2X3R, respectively (Fig. 7 A). Thus, FFA with an IC_50_ value of 221.7 µM was less potent than diclofenac in hP2X3R inhibition. In contrast to diclofenac, FFA did inhibit rP2X3R-mediated currents, although it was more potent at the hP2X3R than at rP2X3R (IC_50_ value of 221.7 µM and 264,1 µM, respectively). This indicates a weaker selectivity of FFA towards the human P2X3R in comparison to diclofenac. Importantly, a concentration of 100 μM FFA, which is commonly used in research to avoid bias resulting from the activation of various other ion channels, exerted a relevant inhibitory effect of 30 % on hP2X3R and 25 % on rP2X3R (Fig. 7 A).

**Figure 7.**
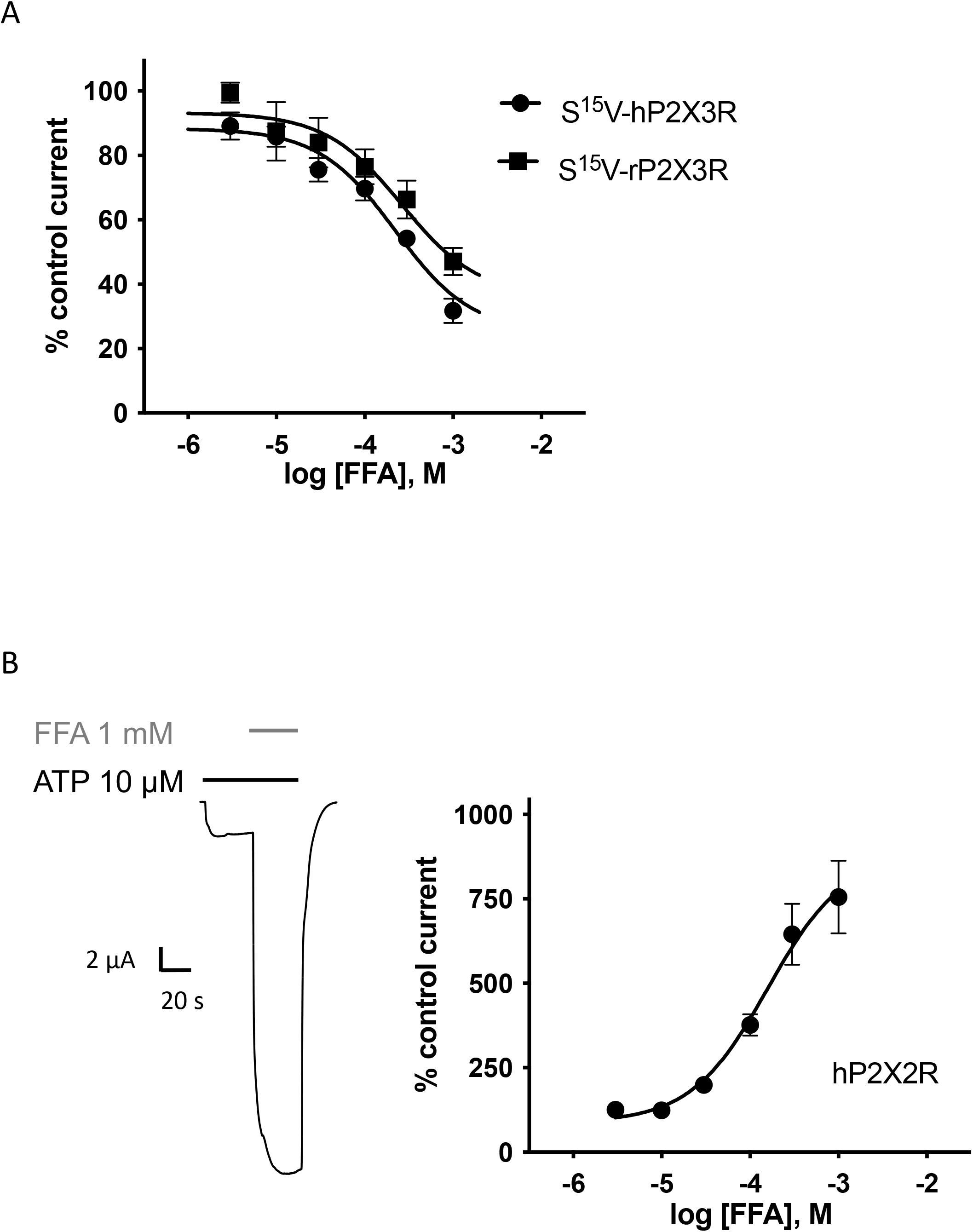
FFA inhibits P2X3R-mediated responses but potentiates hP2X2R-mediated responses. (A) Concentration–inhibition curve of FFA at the human (●) and rat (_□□_) S^15^V-P2X3R exhibited IC_50_ values of 221.7 µM (95% CI: 98.9 - 497 µM) and 264.1 µM (95% CI: 56.9 - 612 µM), respectively. Data points represent the means and SEM. (B) Representative original current trace shows the effect of 1 mM diclofenac (grey bar) on the ATP-induced (10 μM, black bar) current mediated by the hP2X2R expressed in *X. laevis* oocytes. (C) Concentration–response curve of FFA at the hP2X2R (●) exhibited half maximal potentiation value of 43.9 µM (95% CI: 10.6 - 182.5 µM). Data points represent the means and SEM.

In case of FFA, selectivity profiling was performed at hP2X2R and hP2X7R. These two subtypes were chosen, because these either were potentiated or are often analyzed in presence of FFA, respectively. When ATP and FFA were coapplied at the hP2X2R, the current amplitude increased up to 8-fold compared to the steady state current in absence of FFA (Fig. 7 B). Thus, in comparison to diclofenac, FFA shows a significantly higher potentiating effect on hP2X2Rs. The effect of FFA on hP2X7R-mediated currents was assessed by applying concentrations of 100, 300 and 1000µM. A concentration of 100μM, which is commonly used in research applications as a supplement to divalent free ORi-for recordings of the P2X7R, exerted a relevant inhibitory effect of ∼ 39 % (Suppl. Fig. 5). A rough assessment of the IC_50_ value using the three tested concentrations suggested a value of approximately 900 μM for hP2X7R inhibition. These results demonstrate that the use of 100 µM FFA in the analysis of hP2X2R, hP2X3R, and hP2X7 receptors must be critically evaluated, because FFA significantly modulates receptor function, which may result in a significant bias if receptor function is to be quantified.

## 4 Discussion

### 4.1 Inhibition of hP2X3R-mediated currents as an additional mode of action of NSAIDs

Our findings demonstrate the inhibition of the human P2X3R by various NSAIDs. Diclofenac proved to be the most effective antagonist with an IC_50_ value of 138.2 µM. Among the investigated NSAIDs, diclofenac, FFA and flunixin exerted an enduring, potentially irreversible inhibitory effect on the current amplitude, which could not be eliminated by the following washout period.

Considering the involvement of hP2X3R in nociception, it is conceivable that inhibition of hP2X3R contributes to the analgesic effect of NSAIDs and represents an additional mode of action besides COX inhibition. However, plasma levels and IC_50_ values must be taken into consideration. In case of diclofenac, low nanomolar plasma levels are reached during transdermal application, whereas 10-20-fold higher concentrations can be observed in synovial tissue (Efe et al., 2014). When injected intramuscularly, significantly higher plasma levels of approximately 1.8 μg/ml (∼ 6 μM) can be achieved (Drago et al., 2017). Similarly, maximum plasma levels of approximately 2.3 - 2.6 µg/ml (∼ 7 - 8 μM) can be achieved with oral application of 50 - 75 mg diclofenac (Kienzler et al., 2010, Kurowski et al., 1994). According to our experimental data, the current amplitude of the hP2X2/3R and hP2X3R were reduced by approximately 20-30% in presence of 3-10 µM diclofenac. Therefore, it can be assumed that clinically relevant concentrations of diclofenac exert a significant inhibitory effect on hP2X3R-mediated currents. However, diclofenac shows an manyfold higher potency at COX1 (IC_50_ value of 0.075 µM) and COX2 (IC_50_ value of 0.038 µM) than at hP2X3R (IC_50_ value of 138.2 µM) (Warner et al., 1999). The effect of diclofenac besides COX-inhibition has also been studied by other groups, such as (Gan, 2010). For instance, in addition to COX inhibition, other effects such as inhibition of acid-sensing ion channels (ASICs) were discovered (Voilley et al., 2001). However, inhibition of P2X3R has not yet been described before.

Similar to our findings for diclofenac, Hautaniemi et al. described the inhibition of P2X3R by the NSAID naproxen (Hautaniemi et al., 2012). According to our TEVC data, high micromolar to low millimolar concentrations of naproxen are necessary to inhibit hP2X3R, indicating a low potency of naproxen. These results are consistent with the results that Hautaniemi et al. obtained from calcium imaging of rat trigeminal neurons (Hautaniemi et al., 2012). Despite its lower potency in comparison to diclofenac, naproxen might as well exert a relevant inhibitory effect due to higher plasma levels. When administered orally, maximum plasma levels of approximately 70 - 80 µg/ml (∼ 300 - 350 µM) are reached after about two hours (Desager et al., 1976, Dresse et al., 1978).

### 4.2 Selectivity profiling of diclofenac at P2X receptors and related side effects

Selectivity profiling of diclofenac at the different P2XR subtypes showed a strong inhibition of hP2X3R and hP2X2/3R and a weaker inhibition of hP2X1R, hP2X4R and hP2X7R. The rank order of its potencies is: hP2X2/3R > hP2X3R > hP2X1R > hP2X4R, hP2X7R.

Diclofenac had a similar maximum inhibitory effect on hP2X1R as on hP2X3R, but a lower potency. In contrast to its potentially irreversible effect on hP2X3R, diclofenac seemed to exert a reversible inhibitory effect on hP2X1R. Inhibition of hP2X1R, which is involved in inflammatory responses (Lecut et al., 2009), might contribute to the anti-inflammatory effect of diclofenac. However, the difference in potency between hP2X1R (IC_50_ 337.8 μM) and COX (IC_50_ value of 0.075 µM or 0.038 µM of COX1 or COX2, respectively) should be kept in mind.

Considering its low potency at hP2X4R and hP2X7R, it is unlikely that inhibition of these P2X subtypes, which are involved in nociception (Chessell et al., 2005, Tsuda et al., 2009), contributes to the analgesic effect of diclofenac.

Remarkably, diclofenac had a weak potentiating effect on hP2X2R-mediated currents. Regarding its strong inhibitory effect on hP2X3R and its weak potentiating effect on hP2X2R, a predominantly inhibitory effect on the heterotrimeric hP2X2/3 receptor could have been expected. Surprisingly, diclofenac even seemed to be more potent at hP2X2/3R (IC_50_ 76.7 µM) than at hP2X3R (IC_50_ 138.2 µM). However, it should be kept in mind that the use of the correction factor in hP2X3R measurements and the use of different agonists at hP2X2/3 and hP2X3R implies a bias that may affect an accurate, direct quantitative comparison. Since the run down varies between different recordings, the inhibitory effect of diclofenac on hP2X3R might be underestimated.

It is presumed that taste disorders, which have been observed in clinical trials of newly developed P2X3R antagonists, mainly result from the inhibition of heterotrimeric P2X2/3R (Garceau and Chauret, 2019, McGarvey et al., 2022). Therefore, the question arises whether diclofenac might cause taste disorders due to its inhibitory effect on hP2X2/3R. While taste disorders as a side effect are listed as “very rare” in the prescribing information, more than 110 suspected cases of ageusia, dysgeusia or taste disorders have been reported in the European Union so far in EudraVigilance (up to 28/11/2022) for diclofenac (http://www.adrreports.eu/de/). It must also be assumed that there is a high number of unreported cases. In a prospective, randomized clinical trial regarding postoperative administration of Diclofenac, 58% of patients reported impaired taste sensation (Attri et al., 2015). Therefore, it seems likely that diclofenac affects taste sensation due to the inhibition of hP2X2/3R.

### 4.3 Mechanism of action of diclofenac at P2X3R and functional implications for gating

We have found the following lines of experimental evidence for a competitive inhibition of P2X3-subunit containing receptors by diclofenac: (i) inhibition by 30 µM diclofenac of the hP2X2/3R could be overcome by increasing concentrations of the agonist α,β-meATP; (ii) at the L^191^A/S^15^V-hP2X3R the inhibition by diclofenac or FFA at the beginning of the co-application with ATP could be overcome by prolonged ATP co-application; (iii) the L^191^F, N^190^A and the G^189^R mutants of the negative allosteric binding site (Obrecht et al., 2019, Wang et al., 2018), which markedly affected the extent of inhibition of the hP2X3R by the allosteric antagonists gefapixant/AF-219 or ATA were inhibited by 100 µM diclofenac to a similar extent than the hP2X3R.

In addition, our extensive all atom molecular dynamics simulation studies have determined that the most common binding pose of diclofenac at the hP2X3R largely overlaps with ATP bound to the open-state conformation of the hP2X3R. Furthermore, we show by RMSF analysis that diclofenac when bound to the hP2X3R restricts the conformational flexibility of the left flipper and dorsal fin domains, crucially implicated in ATP-induced gating of the hP2X3R (Mansoor, 2022, Mansoor et al., 2016). Our molecular dynamics simulation results also provide a mechanistic explanation for the inhibition of the ATP-induced gating of the hP2X3R: the strong interactions of diclofenac with the residues K201 and E270 of the dorsal fin and left flipper domains, respectively, prevent the conformational rearrangements of dorsal fin and left flipper domains. These are otherwise essential for channel gating or mechanistically, for the transmission of ATP binding to the conformational rearrangement of the lower body domain and eventually opening of the ion channel pore. Thus, our results show that diclofenac acts via a similar mechanism of action than TNP-ATP (Mansoor et al., 2016).

### 4.4 Selectivity profiling of FFA at P2XRs questions its use in P2XR assays

FFA proved to inhibit human P2X3R-mediated currents with a lower potency (IC_50_ value of 221.7 µM) than diclofenac (IC_50_ 138.2µM). In contrast to diclofenac, FFA did inhibit rat P2X3R-mediated currents (IC_50_ value of 264,1µM), indicating a weaker selectivity of FFA towards the human P2X3R. As FFA is usually applied transdermally, plasma levels do not exceed 180 nM even with repetitive application (Drago et al., 2017). Due to its low potency at the hP2X3R and its low plasma levels, it must be assumed that P2X3R inhibition does not contribute to the analgesic effect of FFA in a relevant manner.

However, the inhibitory effect of FFA on P2X3R could be of importance for other scientist that perform functional recordings of P2XRs with solutions supplemented with FFA. Being a non-selective ion channel blocker, FFA is widely used in research to avoid bias resulting from the activation of various other ion channels (Guinamard et al., 2013). Our findings demonstrate that a commonly used concentration of 100 μM FFA exerts a relevant inhibitory effect of approximately 30% on hP2X3R and 25% on rP2X3R. However, especially for recordings of the P2X7R, FFA is often used as a supplement to divalent free ORi-solution (Hülsmann et al., 2003). While inhibition of the P2X3R by FFA has not yet been described by other groups, there are contradictory results regarding its effect on P2X7R. Suadicani et al. suggested a competitive antagonism of FFA at P2X7R (Suadicani et al., 2006), whereas Ma et al. did not find an inhibitory effect of FFA on P2X7R, but rather an inhibition of pannexin-1 by FFA (Ma et al., 2009). According to our data, hP2X7R-mediated currents are reduced by approximately 40% in presence of 100 μM FFA. Even when FFA is no longer applied, the permeability of the receptor remains affected as shown in Suppl. Fig. 5 (Suppl. Fig. 5) by comparing the slope of the linearly increasing current before and after FFA application, which may indicate a potentially irreversible effect of FFA on hP2X7R-mediated current responses.

Taken together, our results question the use of FFA as a nonselective ion channel blocker when P2XR-mediated currents are to be measured.

In comparison to diclofenac, FFA shows a significantly higher potentiating effect on hP2X2R with a 7-8-fold increase in current amplitudes. This potentiating effect of FFA on hP2X2R has already been described by other groups (Schmidt et al., 2018). However, Schmidt et al. do not attribute this effect to a direct interaction of FFA with the receptor, but to membrane alterations caused by the amphiphilic FFA (Schmidt et al., 2018). However, from our point of view, this theory seems questionable, since it cannot explain the opposing effect of FFA on hP2X2R and hP2X3R.

## 5 Conclusion/Summary

In a previous screening of 2000 approved drugs, natural products and bioactive substances, various NSAIDs were found to inhibit S15V-rP2X3R-mediated currents (Obrecht et al., 2019). Using TEVC, we identified diclofenac as a hP2X3R and hP2X2/3R antagonist with micromolar potency (with IC_50_ values of 138.2µM and 76.72µM, respectively), which also showed to be effective in antagonizing native P2X2/3R-mediated responses in pig DRG neurons. Considering their involvement in nociception, inhibition of hP2X3R and hP2X2/3R by micromolar concentrations of diclofenac may contribute to the analgesic effect of diclofenac and represent an additional -although less potent-mode of action besides the well-known COX inhibition. Our results strongly support a competitive antagonism through which diclofenac, by interacting with residues of the ATP-binding site, left flipper, and dorsal fin domains inhibits gating of P2X3R by conformational fixation of the left flipper and dorsal fin domains. A less potent inhibition of hP2X3R was observed for all other investigated NSAIDs. FFA proved to inhibit hP2X3R, rP2X3R and hP2X7R, questioning its use as a non-selective ion channel blocker, when P2XR-mediated responses are under study.

## Contribution to the field statement

P2X3 receptors (P2X3R) are ion channels of sensory neurons and are involved in nociception. P2X3R inhibition reduces neuropathic and chronic pain as well as chronic cough and taste sensations. A previous study suggested that several nonsteroidal anti-inflammatory drugs (NSAIDs) inhibit P2X3R. Here, we examined several NSAIDs at P2X3R and other related P2XR and identified diclofenac for the first time as an hP2X3R and hP2X2/3R antagonist with micromolar potency. Another NSAID, flufenamic acid proved to be an inhibitor of P2X3R and hP2X7R, questioning its widespread use as a nonselective ion channel blocker in P2XR assays.

Pharmacological experiments suggest a competitive mechanism of antagonism of diclofenac that was further supported by molecular dynamics simulations, which demonstrated that diclofenac interacts with the ATP binding site of the hP2X3R. The inhibition is caused by restricting the conformational flexibility of the left flipper and dorsal fin domains of P2X3R, which usually is crucial for ATP-induced channel opening.

Inhibition of hP2X3R and hP2X2/3R by micromolar concentrations of diclofenac may contribute to both the analgesic effect and the side effect of taste disturbance of diclofenac and may represent an additional mechanism of action besides the highly potent COX inhibition.

## 9 Author Contributions

LG, AO, GS, JM, AL, and RH were involved in study design; LG, LC, SC, BH, IT, JK, LE, JM, AL, and RH were involved in data collection and interpretation. LG wrote the first draft of the manuscript. All authors contributed to manuscript revision, read, and approved the submitted version.

## 11 Acknowledgments

The authors gratefully acknowledge the computing time granted through JARA on the supercomputer JURECA at Forschungszentrum Jülich and the supercomputer CLAIX at RWTH Aachen University.

## 12 Supplementary Material

### 12.1 Supplementary Figures and Tables

Supplementary Figures are available under the following link …….

## 13 Data Availability Statement

Relevant materials such as study material (e.g. cDNA of oocyte expression plasmids of the P2XR under study) or individual datasets are available upon request to interested researchers. If desired, the corresponding author Ralf Hausmann, RWTH Aachen University, Germany should be contacted via email.

