## Supplementary material for "Diclofenac and other Non-Steroidal Anti-Inflammatory Drugs (NSAIDs) are Competitive Antagonists of the human P2X3 Receptor": Suppl. Material

### 1 Supplementary Figures and Tables

#### 1.1 Supplementary Figures

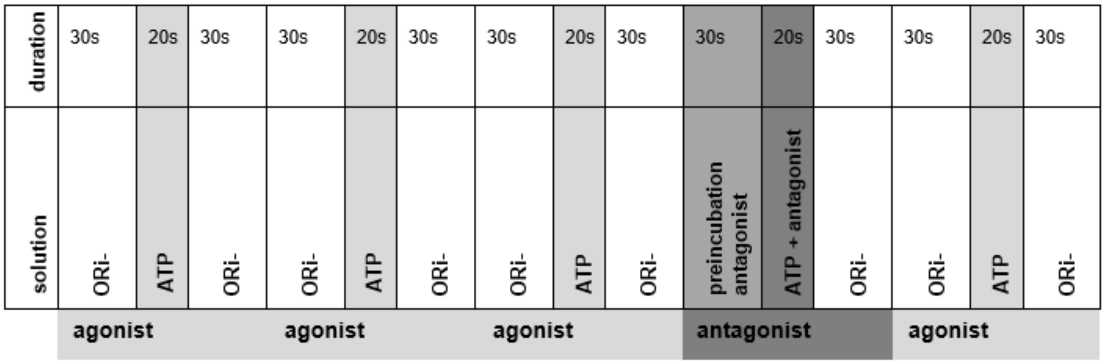


**Supplementary Figure 1.** Peak current protocol used for recordings of the desensitizing P2X1R and P2X3R mutants. First, the agonist ATP was applied three times for 20 seconds each to obtain a stable current amplitude as a reliable reference value. Each ATP application was followed by a wash out step with ORi- solution for 30 seconds. The wash out step was extended to 60 seconds to stabilize the baseline if necessary. After three applications of ATP and a washout step with ORi- solution, a 30-second preincubation with the antagonist and the simultaneous application of ATP and the antagonist for 20 seconds followed. Finally, ATP was once more administered in the absence of the antagonist.

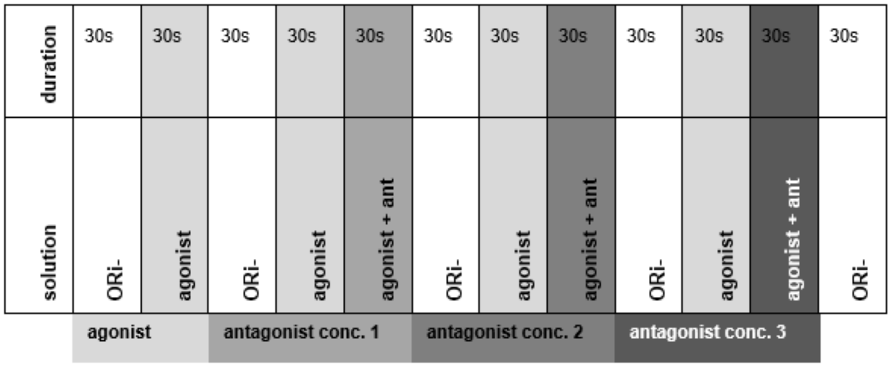


**Supplementary Figure 2.** Steady-state protocol used for recordings of the non- or partially desensitizing hP2X2R, hP2X2/3R or hP2X4R. In case of hP2X2R and hP2X4R the agonist ATP was applied to evoke current responses, whereas its derivate α,β-meATP was used for the heteromeric hP2X2/3R. When the agonist was applied, a steady state occurred after a few seconds, i.e. the current amplitude remained constant. After reaching the steady state, agonist and antagonist (ant) were co-applied. After a wash out step with Ori- this procedure could be repeated for further concentrations (conc.) of the antagonist. The wash out step with ORi- was extended to 40 seconds to stabilize the baseline if necessary.

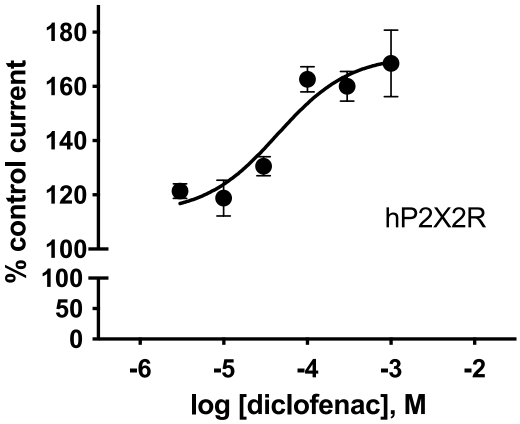


**Supplementary Figure 3.** Diclofenac potentiates hP2X2R-mediated responses. Concentration–response curve of diclofenac at the hP2X2R (●) exhibited half maximal potentiation value of 158.4 µM (95% CI: 64.4 - 389.7 µM). Data points represent the means and SEM.

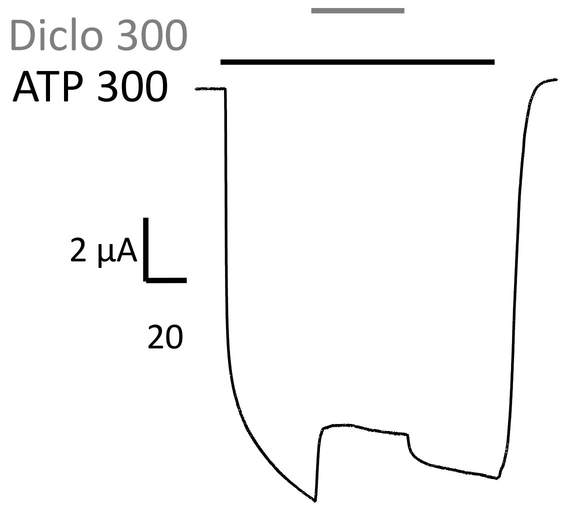


**Supplementary Figure 4.** Effect of diclofenac at hP2X7R. Representative original current trace shows the effects of 300 µM diclofenac (grey bars) on $ATP^{4-}$-induced (300 µM, black bar) currents through hP2X7R expressed in *X. laevis* oocytes.

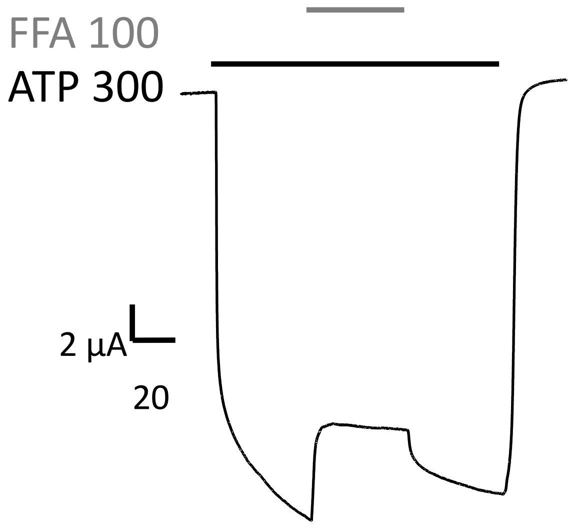


**Supplementary Figure 5.** Effect of FFA at hP2X7R. Representative original current trace shows the effects of 100 µM FFA (grey bars) on $ATP^{4-}$-induced (300 µM, black bar) currents through hP2X7R expressed in *X. laevis* oocytes.

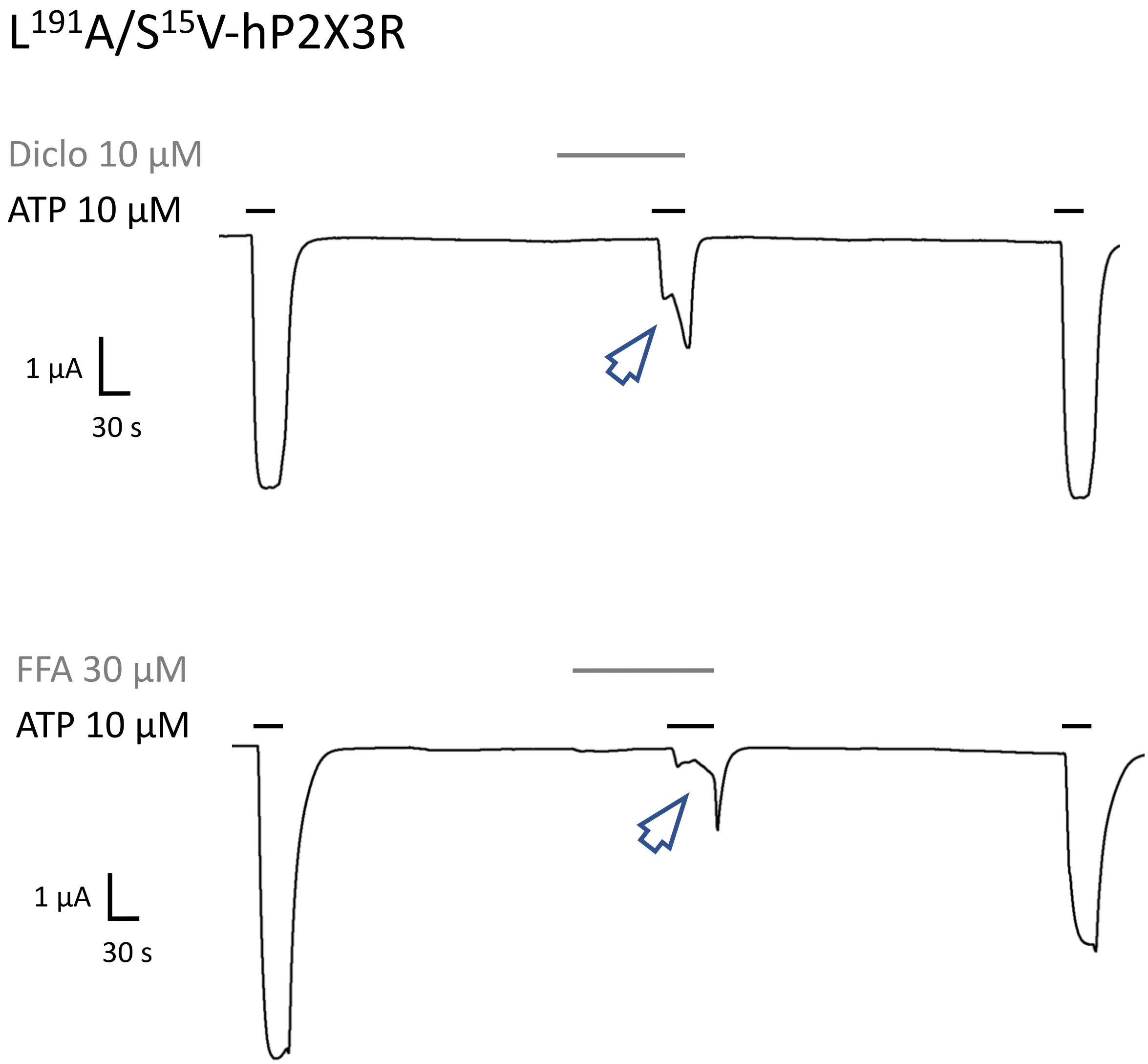


**Suppl. Figure 6.** Effect of diclofenac and FFA at the L$^{191}$A/S$^{15}$V-hP2X3R. Representative original current traces shows the effects of 10 µM diclofenac (upper panel, grey bar) or 30 µM FFA (lower panel, grey bars) on ATP-induced (10 µM, black bars) currents (3$^{rd}$, 4$^{th}$ and 5$^{th}$ ATP application of the peak current protocol are shown). Please note that the initial inhibition during co-application was overcome by prolonged ATP co-application (arrowheads).

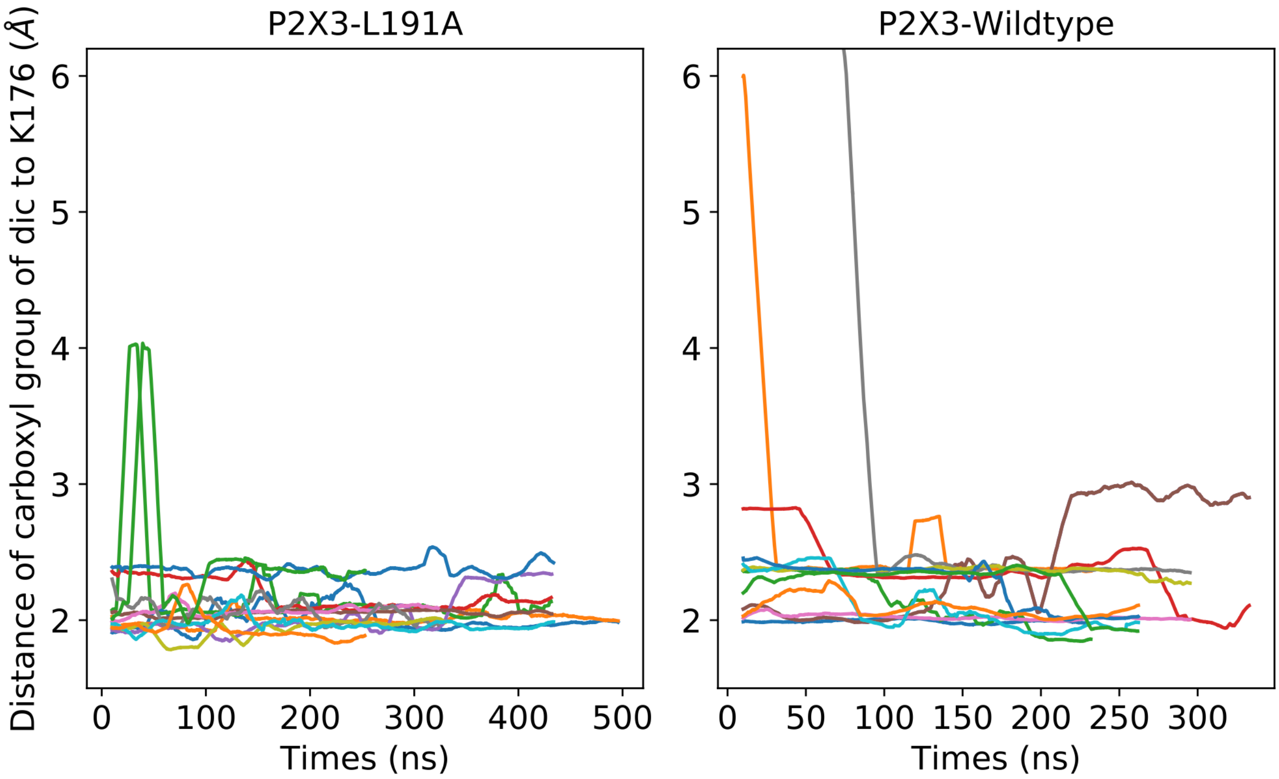


**Suppl. Figure 7.** Distances between carboxyl group of diclofenac to $K^{176}$ shown for all trajectories that were used for clustering for both P2X3 wildtype and $L^{191}A$ mutant over time.

### 1.2 Supplementary Tables

#### Supplementary Table 1

Suppl. Table 1: Volumes and amounts of cRNA injected per oocyte for the different P2XR subtypes.

For the electrophysiologic recordings after 48 hours, a smaller amount of RNA was injected than for the recordings after 24 hours and the oocytes were stored overnight at 4°C instead of 19°C to achieve a similar expression on both recording days. To express the heteromeric hP2X2/3R, the cRNA of His-hP2X2R and wt-hP2X3R were coinjected. For the recordings of the P2X7 receptor, two different constructs (His-hP2X7 and wt-hP2X7) were tested.

| **cRNA** | **Injected volume per oocyte [nl]** | **Injected amount of RNA per oocyte [ng]** |
|---|---|---|
| His-$S^{15}$V-hP2X3R | 41 | Recording after 24 hours: 3.7-4.9<br>Recording after 48 hours: 1.9-2.5 |
| $S^{15}$V-rP2X3R | 41 | Recording after 24 hours: 3.7-3.9<br>Recording after 48 hours: 1.8-1.9 |
| hP2X2/3R (co-injection of His-hP2X$2_A$R and hP2X3R) | 41 | Recording after 24 hours: 2.2 (His-hP2X$2_A$R) + 13.9 (hP2X3R)<br><br>Recording after 48 hours: 0.9 (His-hP2X$2_A$R) + 7.0 (hP2X3R) |
| His-hP2X$2_A$R | 23 | Recording after 48 hours: 0.1 |
| His-hP2X4R | 41 | Recording after 48 hours: 48.0 |
| His-$^{20}$RMVL$^{23}$KVIV$^{23}$,$S^{26}$N-hP2X1R | 23 | Recording after 48 hours: 4.1 |
| His-hP2X7R | 41 | Recording after 24 hours: 1.0 |
| wt-hP2X7R | 41 | Recording after 24 hours: 4.8 |
